# Oxidative stress markers have low repeatability: A meta-analysis and simulation study with implications for measuring physiological condition and fitness

**DOI:** 10.1101/2025.09.02.673724

**Authors:** Rachel R Reid, Davide M Dominoni, Jelle Boonekamp

## Abstract

Markers of oxidative stress are widely used as indicators of physiological state, but their utility for ecological and evolutionary inference remains uncertain due to high intraindividual variation masking associations with environmental conditions, fitness, and other physiological indicators such as telomere length. Although numerous longitudinal studies exist, individual repeatability of oxidative stress measurements is rarely reported explicitly. Here, we present the first meta-analysis assessing individual repeatability of oxidative stress markers, comprising 123 repeatability estimates from 22 studies. Overall, oxidative stress markers exhibited low repeatability on average (Intraclass correlation = 0.164), although repeatability estimates were highly heterogenous across biomarkers, study systems, and ecological contexts. Repeatability was generally low across taxa, sexes, study designs, and environments, although some markers, particularly lipid peroxidation quantified with HPLC, exhibited moderate repeatability in specific contexts. To illustrate the statistical consequences of low repeatability, we used heuristic simulations examining associations between oxidative stress and telomere length under different biological scenarios. These simulations demonstrate how low repeatability substantially reduces statistical power, but this is somewhat mitigated by taking repeated measurements. More broadly, our findings highlight the value of greater cross-disciplinary dialogue, as ecological and biomedical studies often use oxidative stress biomarkers in different ways and may benefit from shared perspectives.

## Introduction

Oxidative stress is a pervasive feature of life, implicated in cellular senescence, cell proliferation, inflammation, ageing, and disease (Gil Del Valle *et al*. 2015; Kregel & Zhang 2007; Leyane *et al*. 2022; Pole *et al*. 2016; Sies 2020). It arises when prooxidants, exceed the neutralising capacity of antioxidants (Costantini *et al*. 2010; Costantini & Verhulst 2009; Monaghan *et al*. 2009). When oxidative stress becomes deregulated, this imbalance can initiate a cascade of reactions causing DNA damage, alterations to membrane lipids, and damage to other macro-molecules (Itri *et al*. 2014; Sharma *et al*. 2019; Von Zglinicki 2002). At the organismal level, such cellular damage is widely hypothesised to contribute to senescence and ageing-related disease (Campisi 2003; Chen *et al*. 2007; Liguori *et al*. 2018; Luo *et al*. 2020; Pole *et al*. 2016; Reichert & Stier 2017; Speakman *et al*. 2015). Consequently, oxidative stress has become a major focus across ecology, evolutionary biology, veterinary medicine, and biomedicine as a potential proximate cause of variation in physiological state, linking the impacts of environmental stress, to ageing, health, and fitness (Cohen *et al*. 2010; Frijhoff *et al*. 2015; Speakman & Garratt 2014; Vágási *et al*. 2019).

Although oxidative stress markers are widely used across disciplines, their inferential goals differ substantially. In biomedical and veterinary contexts, oxidative stress markers are often interpreted relative to physiological thresholds or disease status, where deviations from a normal physiological range may indicate pathology or disrupted homeostasis (Celi 2011; Dănulescu & Costin 2012; Dhama *et al*. 2019). In contrast, ecological and evolutionary studies frequently use oxidative stress biomarkers to infer differences among individuals in condition, environmental quality, life history trade-offs, or fitness (Boonekamp *et al*. 2017; Smith *et al*. 2016; Vágási *et al*. 2019; Vitousek *et al*. 2016). Many ecological applications implicitly assume that oxidative stress measurements capture biologically meaningful and consistent variation among individuals across time, *in the healthy population*. This notably differs from the clinical contexts in which abnormal oxidative stress profiles are typically considered in association with pathology. Instead, ecological interest in oxidative balance has largely centred on the idea that individuals with below average oxidative balance may experience reduced survival, reproduction, or other fitness costs (Speakman *et al*. 2015; Speakman & Garratt 2014; Vágási *et al*. 2019). However, studies investigating oxidative stress and fitness-related traits have often yielded heterogeneous results, suggesting that the interpretation of oxidative stress biomarkers may depend strongly on environmental, methodological, and biological context (Speakman & Garratt 2014; Vitousek *et al*. 2016).

One potential explanation for the observed heterogeneity amongst study findings is that oxidative stress physiology is highly dynamic, and measurable markers of oxidative stress may therefore exhibit high intraindividual variability within relatively short timescales. These oxidative stress dynamics may strongly depend on life stage and environmental context and therefore be poorly predictable and generalisable. If oxidative stress fluctuates unpredictably through time, then observed measurements may poorly reflect persistent differences among individuals across environmental contexts. Such variation presents a particular challenge in ecological and evolutionary research as high within-individual variability may obscure associations between physiology, environment, and fitness, even when repeated measurements are collected. Importantly, low repeatability does not necessarily invalidate the biological relevance of a biomarker, nor does it preclude its usefulness for detecting acute physiological changes or pathological conditions. Rather, low repeatability limits the extent to which oxidative stress measurements can be used for statistical inference about persistent individual differences in condition, physiology or fitness, and in particular in the situation when the observed intra-individual variation is both large and unpredictable (Fig. 1). Conceptually, the utility of oxidative stress biomarkers therefore depends not only on the magnitude of intraindividual variation, but also whether that variation is predictable or can be statistically accounted for (Fig 1).

**Figure 1.**
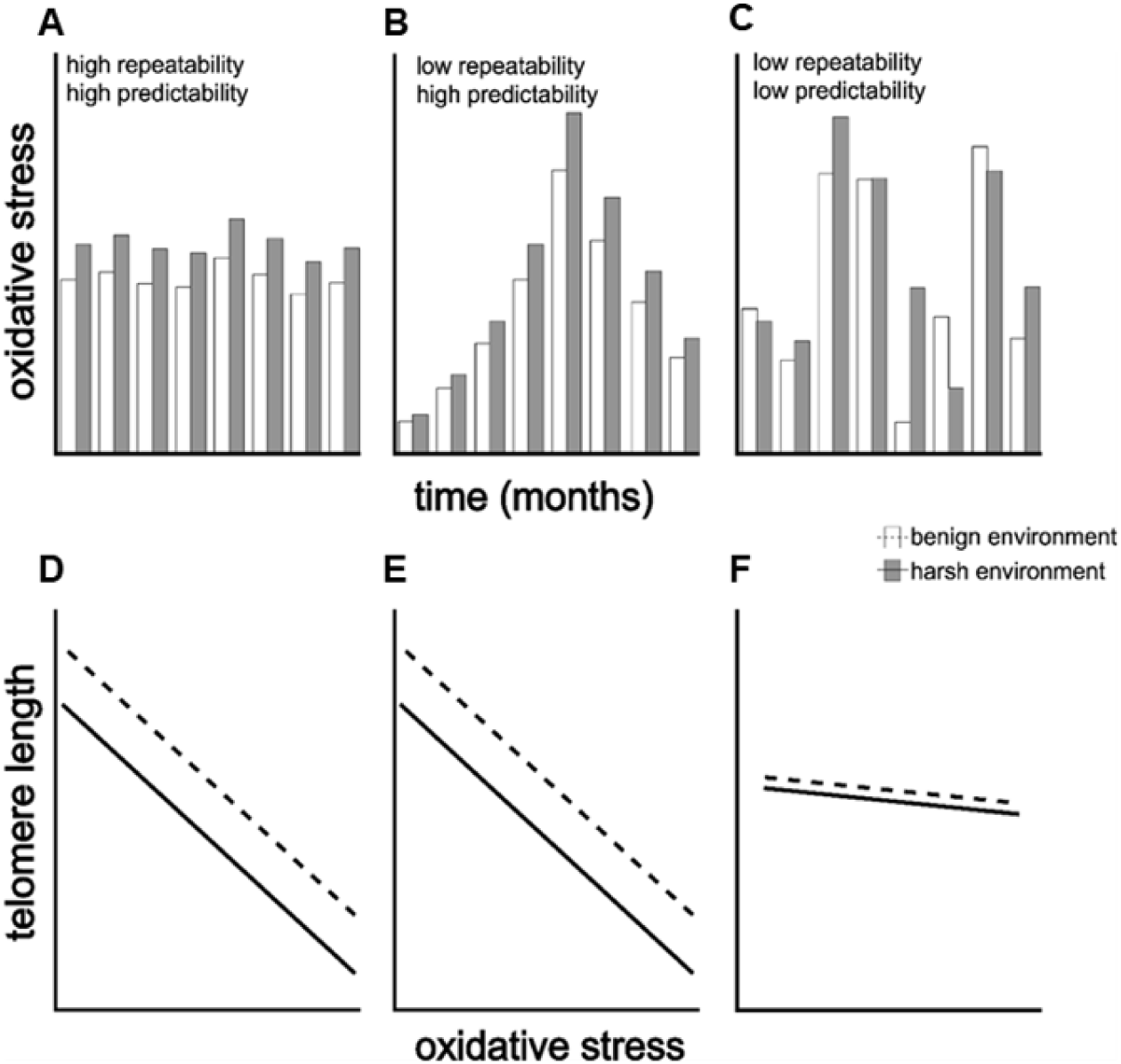
Conceptual depiction of how different forms of intraindividual variation influence the utility of oxidative stress measurements as biomarkers of environmental conditions and somatic state, here conceptualised as telomere length. Panels (A-C), depict repeated individual measurements of oxidative stress across time for (A) high repeatability and predictability, (B) low repeatability and high predictability, (C) low repeatability and low predictability for an individual living in a benign (white bars), and another individual living a harsh environment (grey bars). Panels (D), (E), (F), illustrate the impact of these corresponding scenarios on the population-level associations between oxidative stress, telomere length, and the two environmental conditions. (E) illustrates that there is little negative impact of low repeatability when one can statistically control for the confounding variation. In contrast, panel (F) illustrates how high and unpredictable intraindividual variation can obscure associations between oxidative stress, environmental conditions, and physiological traits such as telomere length.

It is important to make the distinction between repeatability, biomarker utility, and the true physiological causes and consequences of oxidative stress. Ecological studies often use oxidative stress biomarkers to understand variation in condition, health, and fitness, yet relationships with fitness-related traits are frequently inconsistent and highly context dependent (Beaulieu & Costantini 2014; Cohen *et al*. 2010; Speakman & Garratt 2014). Weak correlations between markers may also arise when oxidative stress markers exhibit high intraindividual variability, even if biologically important causal relationships exist. One illustrative example is the causal role of oxidative stress in telomere attrition. While in vitro studies have shown very pronounced effects of oxidative stress on telomere attrition (Von Zglinicki 2002), this relationship is virtually absent in vivo (r = 0.027) as observed in a meta-analysis of correlational studies (Armstrong & Boonekamp 2023). There is no doubt about that oxidative stress contributes to telomere attrition in vivo, and indeed the same meta-analysis study found stronger support from experimental studies which manipulated oxidative stress (Armstrong & Boonekamp 2023), which illustrates the distinction between the biological importance of a causal effect versus the utility of oxidative stress measurements as biomarker of these processes. Given the widespread use of oxidative stress as biomarkers of physiological condition, it is important to determine how well such measurements distinguish within-from between-individual variation.

For the same reason as outlined above, establishing repeatability has been a key research focus for numerous physiological and behavioural traits, including metabolic rate (Nespolo & Franco 2007), glucocorticoid hormones (Schoenemann & Bonier 2018; Taff *et al*. 2018), cognitive performance (Cauchoix *et al*. 2018), and telomere length (Kärkkäinen *et al*. 2022). However, similar knowledge about the repeatability of oxidative stress markers has not yet been established. Where repeatability estimates have been reported, they vary widely across studies and biomarkers. For example, glutathione peroxidase displayed moderate repeatability (r = 0.47) in great tits (*Parus major)* across seasons (Cláudia Norte *et al*. 2008), whereas reactive oxygen metabolites in collared flycatchers (*Ficedula albicollis)* were effectively non-repeatable (r = 0.03), despite antioxidant defences in the same study showing moderate repeatability (r = 0.581) (Récapet *et al*. 2019). A growing body of evidence suggests oxidative stress is influenced by numerous environmental and life history factors. For example, oxidative stress levels have been shown to vary across life stages (A. Cohen *et al*. 2008; Alonso-Álvarez *et al*. 2010; Monaghan *et al*. 2009), increase during reproduction and mating efforts (Alonso-Álvarez *et al*. 2010; Cram *et al*. 2015), and change during energetically demanding activities such as seasonal migration (Jenni-Eiermann *et al*. 2014). It is therefore unsurprising that oxidative stress physiology varies dynamically in concordance with life-history stages and events, but it remains unclear to what extent such variability is predictable and generalisable, key prerequisites for the utility of oxidative stress measurements as biomarkers.

Further complicating this issue, oxidative stress can fluctuate over very short timescales (A. Cohen *et al*. 2008; Mallard *et al*. 2020), raising concerns about fine-scale temporal stability. In addition, oxidative stress biomarkers encompass a highly diverse family of molecules that differ substantially in their biological function, regulation, and temporal dynamics (Costantini 2019; Costantini *et al*. 2010). Reactive oxygen species (ROS) vary widely in reactivity and half-life, while antioxidant components can be either enzymatic or chemically derived through dietary factors, with vastly different specificity toward ROS as well as localization (Murphy *et al*. 2022). Likewise, biomarkers such as, antioxidant enzymes, non-enzymatic antioxidants, and measures of oxidative damage may reflect distinct physiological processes rather than a single unified oxidative state. This biochemical diversity may help explain why different oxidative stress markers are often weakly correlated, even within the same individual (Beaulieu & Costantini 2014; Costantini 2019; Costantini *et al*. 2010; Speakman *et al*. 2015). Together, these issues highlight the need to quantify intraindividual variation in commonly used oxidative stress markers and to determine whether some markers exhibit greater repeatability, and therefore potentially greater utility for ecological and evolutionary inference, than others. Furthermore, there is a need to determine if common denominators exist that may help explaining and generalising the observed intraindividual variation as this would further increase the utility of oxidative stress measurements as biomarkers in this context.

The increasing availability of longitudinal datasets now provides the opportunity to formally investigate repeatability of oxidative stress markers across a wide range of non-model species and study systems. Here, we present the first meta-analysis of individual repeatability in oxidative stress biomarkers using non-human datasets. Specifically, we evaluate repeatability in the context of ecological and evolutionary inference by estimating the overall repeatability of oxidative stress markers and identifying potential biological and methodological sources of variation in repeatability. We additionally examine correlations among oxidative stress biomarkers to assess whether consistent multivariate patterns exist across studies. Finally, we use heuristic simulations to illustrate how low repeatability may influence statistical power under different biological scenarios and study designs.

For these simulations we use the example of oxidative stress and telomere attrition. Together, these analyses aim to clarify the extent to which oxidative stress biomarkers capture persistent individual differences and to identify the challenges, and potential solutions, associated with using oxidative stress measurements in ecological and evolutionary research.

## Materials and Methods

### Literature search

We developed a search string using keywords derived from a review of the oxidative stress literature as follows:

**( “Oxidative stress” OR “Antioxidant capacity” OR “Antioxidant*” OR “Oxidative damage”) AND (“Repeated*” OR “Repeated measure” OR “Repeatability*” OR “Longitudinal*” OR “Longitudinal study” OR “Within Individual” OR “Within Individual study”) AND (“Wildlife*” OR “Wild animal” OR “Wild population” OR “Wild*” OR “Bird*” OR “Avian*” OR “Invertebrate*” OR “Fish*” OR “Herpetofauna*” OR “Vertebrate*” OR “Mammal*”) NOT (“Human*” OR “Woman*” OR “Women*” OR “Man*” OR “Children*” OR “Patients*” OR “People*” OR “Men*” OR “Child*”)**

This search string was used to collect literature from both the Web of Science core collection database (last search 7^th^ of December 2023) and Scopus database (last search 4^th^ of June 2024). After removing duplicates, we recorded the number of results for each iteration following the Preferred Reporting Items for Systematic Reviews and Meta-analysis (PRISMA) framework. The PRISMA diagram summarising the selection process is shown in Fig S1. We were able to include effect sizes from 22 studies (Bodey *et al*. 2019; Christensen *et al*. 2016; Cram *et al*. 2015; Eikenaar *et al*. 2020; Gormally *et al*. 2019; Guindre-Parker *et al*. 2013; Guindre-Parker & Rubenstein 2018; Hau *et al*. 2015; Herborn *et al*. 2016; Kolling *et al*. 2022; Marasco *et al*. 2017; Noguera *et al*. 2017; Orquera-Arguero *et al*. 2023; Récapet *et al*. 2019; Rowe *et al*. 2015; Sauerwein *et al*. 2020; Schull *et al*. 2016; Skrip *et al*. 2016; Těšický *et al*. 2021; Velázquez-Cantón *et al*. 2018; Viblanc *et al*. 2018; Vitikainen *et al*. 2016).

### Inclusion criteria

The inclusion criteria were designed to capture longitudinal studies across ecological, evolutionary, and comparative biological systems while excluding clinical and human focused research.

To be included in the meta-analysis, studies had to meet the following criteria:

1. Measured at least one biomarker of oxidative stress (oxidative damage or antioxidant defence).
2. Took at least two repeated oxidative stress measurements from the same individual, with the time interval between measurements reported.
3. Focused on non-human organisms.
4. Provided access to raw individual-level data, either publicly available or obtained via author correspondence.

We included both experimental and observational studies across a range of taxa and environments to maximise ecological and comparative representation.

### Screening process

We used a two-step screening process to identify eligible studies. First, titles and abstracts were screened using the “MetaGear” (Lajeunesse 2016) package in R (R Core Team 2024) to identify studies involving oxidative stress in non-human organisms. Because repeat sampling was not always explicitly stated in abstracts, we next screened the full texts of selected studies, verifying in the methods sections that repeated measurements on the same individuals were conducted.

### Data extraction

From each eligible study, we extracted biological and methodological information, including species, taxonomic class, sample size, number of repeated samples per individual (mean), time between repeated samples (mean), oxidative stress biomarkers measured, environment, and study type (manipulated vs. non-manipulated). Where available, we additionally extracted information on assay validation metrics including intra-assay and inter-assay repeatability or coefficients of variation (CV) (Table S1). For experimental studies, we separated treatment and control groups, and calculated repeatability estimates independently for each group. To ensure comparability across study types, control groups from experimental studies were pooled with observational datasets and classified as non-manipulated, whereas treatment groups were classified as manipulated.

Environment was categorised as wild, captive, wild captive (wild individuals brought into captivity for the purposes of the study), or livestock (Animals used for agricultural purposes, including horses, cattle, and sheep). Biomarkers were grouped according to whether they measured oxidative damage (lipid peroxidation, protein damage, DNA damage, or nonspecific damage) or antioxidant defence (enzymatic or non-enzymatic) (Table S2).

We acknowledge that oxidative stress markers encompass biochemically diverse molecules that differ in function, regulation, and temporal dynamics. However, further subdivision of not feasible for several marker classes because sample sizes were extremely small, limiting statistical power and preventing robust meta-analytic comparisons. Consequently, the biomarker groupings used here represent a balance between biological specificity and sufficient statistical power.

### Repeatability estimation

To calculate within-individual repeatability of oxidative stress measurements, we obtained raw individual-level data from each study. For studies where raw data was not directly available, authors were contacted directly (Table S3). We estimated repeatability by determining the intraclass correlation coefficient (ICC) using ANOVAs based on equations 4 and 5 from Nakagawa & Schielzeth (2010) (Nakagawa & Schielzeth 2010), along with the standard error and 95% confidence intervals.

We selected the ANOVA-based ICC approach for the primary meta-analysis because repeatability estimates in our dataset were frequently very low (<0.05). Under these conditions, linear mixed-effects model (LMM) based repeatability estimates can become upwardly biased near zero due to variance component estimation constraints, whereas ANOVA-based ICC approaches generally provide more conservative and stable estimates of low repeatability (Kärkkäinen *et al*. 2022; Nakagawa & Schielzeth 2010). Similar approaches combining ANOVA-based and LMM-based repeatability estimates have been used in previous repeatability meta-analyses (Cauchoix *et al*. 2018; Kärkkäinen *et al*. 2022; Taff *et al*. 2018).

Negative ICC values were retained in all analyses. Although repeatability itself cannot biologically be negative, negative ICC estimates can arise through sampling variation when within-individual variance exceeds between-individual variance, particularly when true repeatability is close to zero or sample sizes are small (Liljequist *et al*. 2019). Retaining negative estimates avoids upwardly biasing meta-analytic estimates by selectively truncating low values. Such estimates should therefore be interpreted as statistical values consistent with zero repeatability rather than biologically meaningful negative repeatability.

To assess the robustness of the estimation framework, we additionally compared unconditional repeatability estimates derived using both ANOVA-based ICCs and LMM-based repeatability estimates implemented in the rptR package (Fig S2). Although the two approaches partition variance differently and are therefore not directly equivalent, they produced qualitatively similar estimates of repeatability.

For the primary meta-analysis, we calculated separate repeatability estimates within biologically relevant subsets of the data, including sex, oxidative stress biomarker, and study type (manipulated versus non-manipulated). This stratified approach allowed these variables to be incorporated as moderators within the meta-analytic models, enabling us to investigate whether repeatability differed across biological or methodological contexts. In contrast, the conditional repeatability analyses described below used LMM-based approaches that explicitly incorporated covariates as fixed effects within the repeatability estimation itself.

This yielded 123 effect sizes from 22 studies across 22 species published between 2013 and 2023 (species listed in Table S4).

### Oxidative biomarker correlations

For studies measuring multiple oxidative stress biomarkers within the same individuals, we calculated Pearson’s correlation coefficients between biomarker pairs using the cor function in R. Correlations were calculated using repeated individual-level measurements available from each dataset and were subsequently included in a separate meta-analysis to assess the consistency of biomarker co-variation across studies.

### Meta-analysis approach

All analysis were formed in R version 4.3.3 (R Core Team 2024).

Meta-analyses were conducted using a multi-level meta-analysis structure with the metafor package (Viechtbauer 2010). ICC and Pearson’s correlation coefficients were standardised using Fisher’s Z transformation (Nakagawa & Schielzeth 2010) prior to analysis and back-transformed for interpretation in figures and tables. Fisher’s Z transformation is commonly applied to repeatability and correlation estimates in meta-analysis to allow for estimates to be comparable and for appropriate weighting across studies (Holtmann *et al*. 2017; Nakagawa & Schielzeth 2010). Corresponding sampling variances were calculated using the escalc function.

We first estimated the overall repeatability using an intercept-only random effects model. We then tested whether variation in repeatability could be explained by moderators including study type, environment, taxonomic class, oxidative stress biomarker, mean time between measurements (log-transformed), and the average number of samples per individual (log-transformed). Due to the relatively modest sample size and uneven representation across moderator categories, moderators were tested in separate univariate multilevel meta-regression models rather than a single global model. This approach avoided model overparameterization and reduced issues associated with sparse combinations of categorical predictors. Similar univariate moderator approaches have been used in previous repeatability meta-analyses (Cauchoix *et al*. 2018).

Subgroup analyses additionally compared oxidative damage and antioxidant defence categories using biomarker type as a moderator in a separate analysis (Table S2). An additional meta-analysis model was used to explore biomarker correlations using the same multilevel model structure as before, with biomarker pair identity as a categorical moderator.

All models included study ID and observation ID as random effects to account for non-independence arising from multiple effect sizes originating from the same study and residual heterogeneity among effect sizes within studies (Nakagawa *et al*. 2022; Nakagawa & Schielzeth 2010). Total relative heterogeneity and the contribution of each random effect was quantified using the i2_ml function from the OrchaRd package (Nakagawa *et al*. 2023). Although this multilevel structure accounts for substantial shared variance within studies, some residual non-independence may remain due to unmeasured study-level characteristics.

Lastly, we evaluated potential publication bias by fitting two multilevel meta-regression models, with the square root of inverse sampling variance and mean-centred year of publication as moderators to test for small study effects and time lag bias respectively (Fig S3) (Nakagawa *et al*. 2022).

Methods for simulation analyses are provided in S1 text.

### Conditional repeatability comparison

To evaluate whether controlling for common biological covariates altered repeatability estimates, we performed an additional comparison using a subset of studies (n = 8) for which the raw data included information about age, and in some cases, sex and treatment group. For each oxidative stress biomarker in these studies, we estimated repeatability under four model conditions: (1) unadjusted (raw, no covariates), (2) age adjusted, (3) age and sex adjusted, and (4) age and treatment group adjusted.

Repeatability was calculated using the rptR (Nakagawa & Schielzeth 2010; Stoffel *et al*. 2017) package in R, employing an LMM-based estimation approach. This method allows for covariates to be included as fixed effects within the model, providing estimates of conditional repeatability. We then performed a meta-analysis using repeatability model type as a moderator to assess whether controlling for these covariates influenced repeatability estimates. Supplementary comparisons between ANOVA-based and LMM-based repeatability are presented in Fig S2.

## Results

Our meta-analysis synthesised 123 estimates of individual repeatability, estimated using the intraclass correlation (ICC), derived from 22 published datasets. Across all studies, the overall repeatability of oxidative stress markers was low (ICC = 0.164, 95% confidence interval (CI) = [0.093, 0.233], P < 0.001; Table S5, Fig 2), with substantial heterogeneity among effect sizes (I^2^ = 57.4).

**Fig 2:**
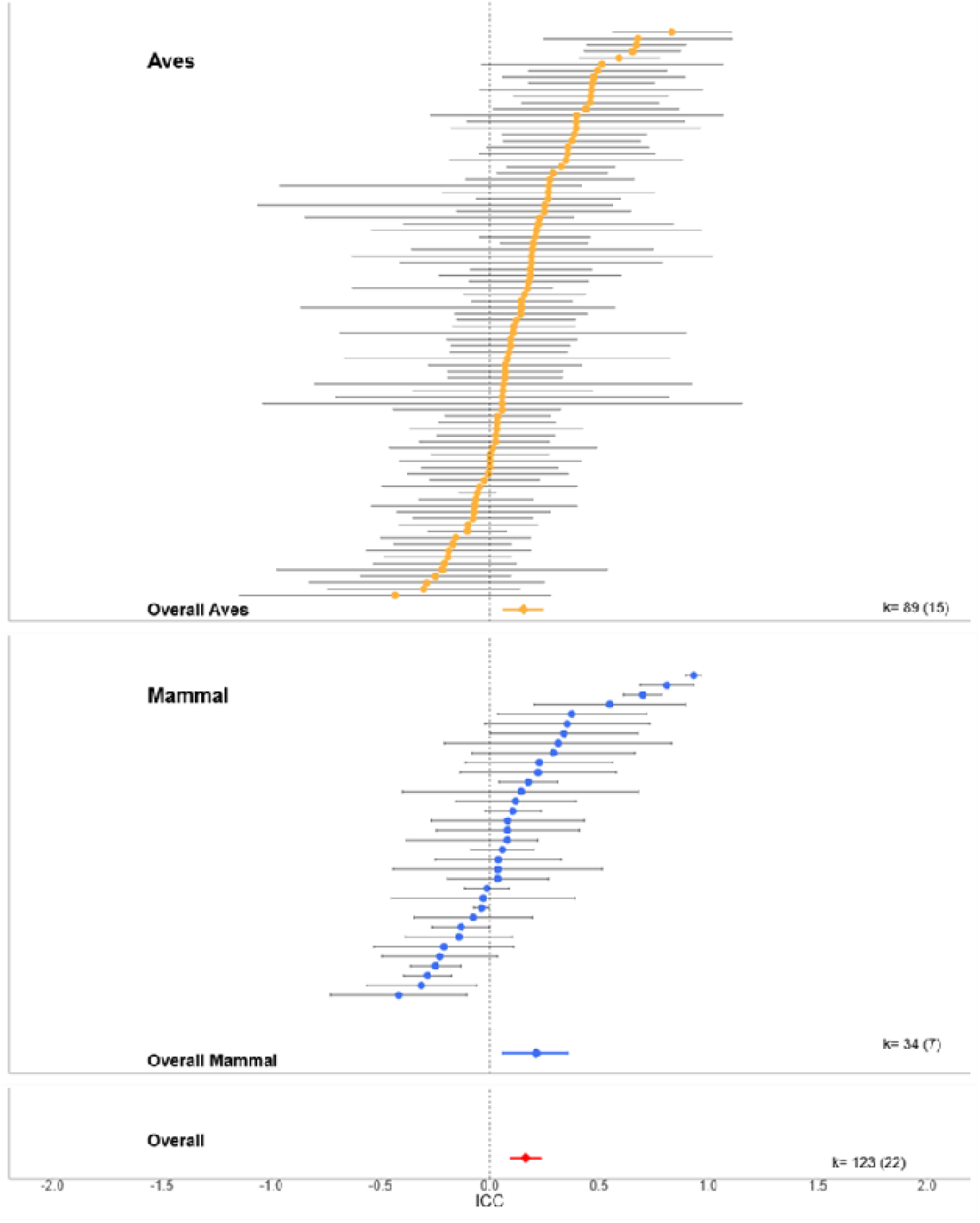
Caterpillar plot displaying individual effect sizes sorted from lowest to highest repeatability and split by taxonomic class. Yellow points represent ICC values for the “Aves” class while blue points correspond to ICC values for the “Mammal” class. The global ICC estimate for “Aves” is shown at the bottom of the Aves class facet, and the global estimate for “Mammals” is shown similary. These estimates were extracted from the meta-model where taxonomic class was included as a moderator. ICC values are presented along with their 95% confidence intervals. The dashed line represents an ICC value of zero. The red diamond and error bars at the bottom of the plot represents the overall ICC value and its 95% confidence intervals from the intercept only model. The corresponding number of effect sizes is shown on the right side of the plot, with the number of studies indicated in parentheses.

Mammals exhibited slightly higher ICC values (ICC = 0.212, 95% CI = [0.058, 0.356], P = 0.007) compared to birds (ICC = 0.155, 95% CI = [0.062, 0.245], P = 0.012; Table S5, Fig 2), but the difference between groups was small and not statistically significant (estimate = 0.059, 95% CI = [-0.123, 0.238], P = 0.526). These findings indicate that oxidative stress markers generally exhibit low repeatability across studies, although substantial heterogeneity among effect sizes suggests that repeatability varies considerably across biomarkers, study systems, and contexts. The broad range of observed ICC values, including instances of moderate to high repeatability, raises the possibility that certain biological or methodological contexts may promote more stable oxidative phenotypes.

### Moderator analyses investigating potential sources of variation in repeatability

We estimated ICC separately within treatment and control groups for experimental studies, and within sex, to enable testing the influence of these variables as moderators in the meta-analysis (see Methods for details). Control groups from experimental studies were pooled with correlational studies to create a two-level moderator comprising manipulated and unmanipulated animals.

Repeatability estimates from unmanipulated animals were slightly lower (ICC = 0.146, 95% CI = [0.053, 0.230]) than those from experimentally manipulated animals (ICC = 0.208, 95% CI = [0.091, 0.319]), although this difference was small and not statistically significant (estimate = -0.064, 95% CI = [-0.204, 0.077], P = 0.86; Table S5, Fig 3A). Similarly, females showed comparable repeatability estimates (ICC = 0.167, 95% CI = [0.065, 0.266]) to males (ICC = 0.154, 95% CI = [0.03, 0.273]; estimate = -0.013, 95% CI = [-0.163, 0.137], P = 0.86; Fig. 3A-B; Table S5, Fig 3B).

**Fig 3:**
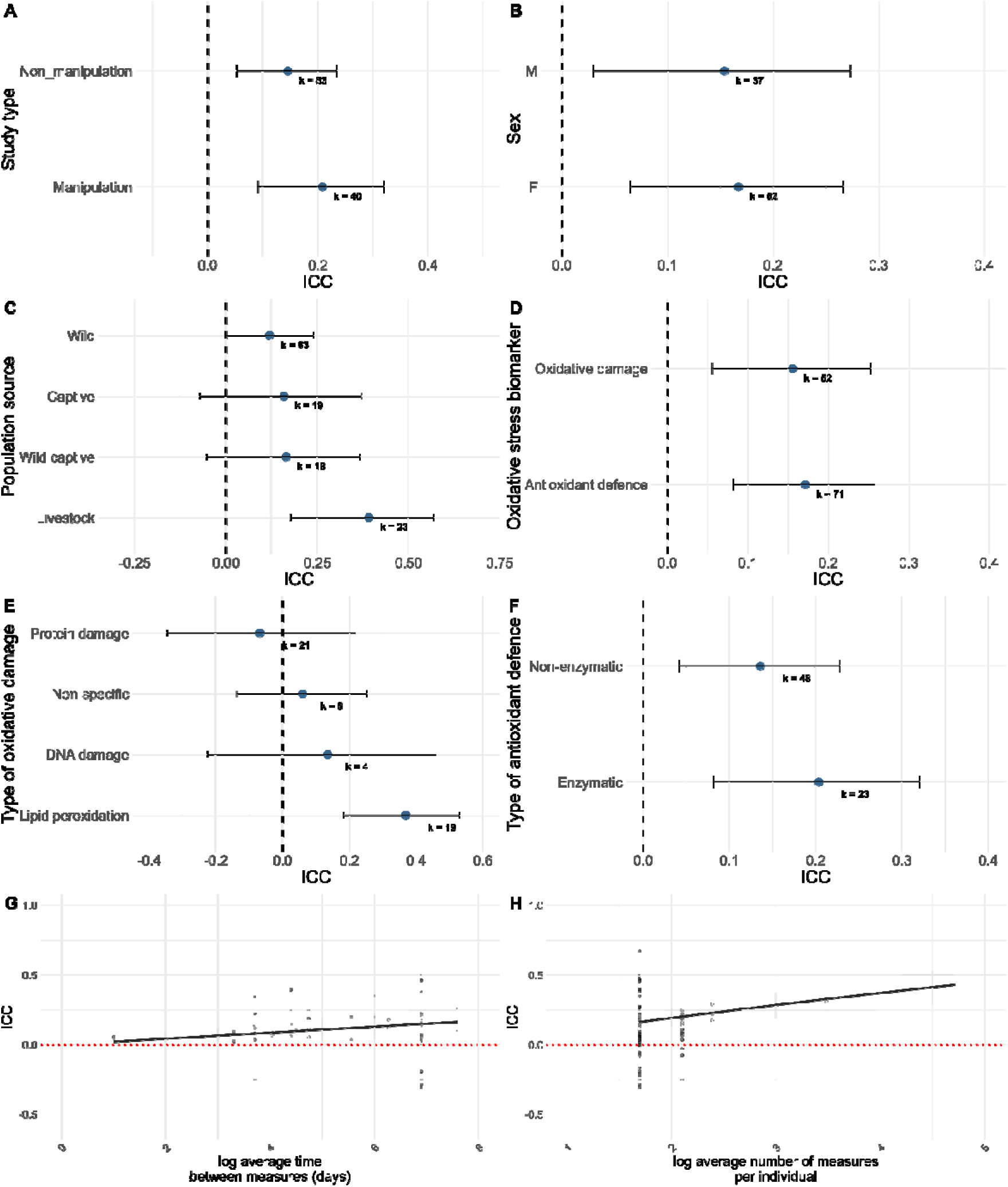
Within-individual repeatability of oxidative stress measures for each moderator level. Panels A-F show repeatability estimates for categorical moderators (A) Study type, (B) Sex, (C) Population source, (D) Oxidative stress biomarker, (E) Type of oxidative damage, (F) Type of antioxidant defence. For each level, the model-estimated repeatability (ICC) is plotted with 95% confidence intervals, back transformed from Fishers Z. The dashed vertical line indicates an ICC of zero. Panels G and H display the relationship between continuous moderators and repeatability: (G) Log-transformed average time between measures and (H) Log-transformed average number of measures per individual. In each panel, grey points represent individual effect sizes, the black line shows the model-predicted relationship, and the shaded ribbon denotes the 95 confidence intervals. The red dashed line indicates an ICC = 0. ICC is plotted on the y-axis in G and H, and on the x axis in A-F.

We observed higher repeatability estimates for livestock populations (ICC = 0.391, 95% CI = [0.178, 0.569], P = 0.005), compared to studies conducted in wild (ICC = 0.119, 95% CI = [-0.003, 0.239], P = 0.057), wild captive (ICC = 0.165, 95% CI = [-0.052, 0.367], P = 0.136), or captive-bred animals (ICC = 0.158, 95% CI = [-0.07, 0.371], P = 0.173; Table S5, Fig 3C). Using livestock as the reference category, model estimates indicated significantly lower repeatability in wild populations (estimate = -0.285, 95% CI = [-0.506, -0.029], P = 0.029). Repeatability was also lower in wild-captive (estimate = -0.242, 95% CI = [-0.513, 0.072], P = 0.129), and captive populations (estimate = -0.248, 95% CI = [-0.523, 0.074], P = 0.074) although neither differed significantly from the livestock reference level. These findings should be interpreted cautiously because livestock datasets generally contain a greater number of repeated measurements per individual, which may contribute to higher ICC estimates.

We found no evidence for systematic differences in repeatability between oxidative damage and antioxidant defence markers (estimate = -0.016, 95% CI = [-0.140, 0.109], P = 0.803). Both categories exhibited low but statistically significant repeatability overall: oxidative damage (ICC = 0.156, 95% CI = [0.056, 0.253], P = 0.002); antioxidant defence (ICC = 0.171, 95% CI = [0.082, 0.258], P = <0.001; Table S5, Fig 3D). Further analysis of oxidative damage marker subtypes revealed that lipid peroxidation markers exhibited comparatively higher repeatability than other oxidative damage marker classes (ICC = 0.368, 95% CI = [0.182, 0.529], P <0.001; Table S6, Fig 3E).

Other damage types, including DNA damage (ICC = 0.135, 95% CI = [-0.224, 0.461], P = 0.464), protein damage (ICC = -0.068, 95% CI = [-0.346, 0.220], P = 0.646), and nonspecific damage (ICC = 0.059, 95% CI = [-0.137, 0.252], P = 0.59), showed little evidence of repeatability. Using lipid peroxidation as a reference level, protein damage (estimate = -0.426, 95% CI = [-0.654, -0.128], P = 0.006) and non-specific damage (estimate = -0.316, 95% CI = [-0.539, -0.051], P = 0.020) exhibited significantly lower repeatability. DNA damage also showed lower repeatability than lipid peroxidation, although this difference was not statistically significant (estimate = -0.245, 95% CI = [-0.578, 0.156], P = 0.228). Enzymatic antioxidants exhibited slightly higher repeatability (ICC = 0.204, 95% CI = [0.081, 0.320], P = 0.001), compared to non-enzymatic antioxidants (ICC = 0.136 95% CI = [0.042, 0.228], P = 0.005; Table S6, Fig 3F), although the difference between these categories was not statistically significant (estimate = -0.069, 95% CI = [-0.198, 0.061], P = 0.296). Both antioxidant classes therefore showed relatively low repeatability overall.

The average time interval between repeated measurements showed a small but statistically significant positive association with ICC (slope estimate = 0.022, 95% CI = [0.006, 0.038], P = 0.008; Table S5, Fig 3G). In addition, the number of repeated measurements per individual showed a significant positive association with ICC estimates (slope estimate = 0.097, 95% CI = [0.064, 0.130], P <0.001; Table S5, Fig 3H).

### Conditional repeatability analysis

To assess whether controlling for individual-level covariates altered repeatability estimates of oxidative stress biomarkers, we conducted a separate meta-analysis using a subset of eight studies for which raw data included information on age, and in some cases sex and treatment group. For each study, we calculated repeatability estimates using four model types: (1) raw repeatability without covariates, (2) age adjusted repeatability, (3) age and sex adjusted repeatability, (4) age and treatment adjusted repeatability. Across all models, repeatability estimates were broadly similar (Fig 4). Raw repeatability estimates (R = 0.195, 95% CI = [0.067, 0.316], P = 0.003), age adjusted repeatability (R = 0.187, 95% CI = [0.058, 0.309], P = 0.004), and age and sex adjusted repeatability (R = 0.167, 95% CI = [0.025, 0.302], P = 0.021) were all statistically indistinguishable. Repeatability estimates adjusting for age and treatment group were somewhat higher (R = 0.356, 95% CI = [-0.035, 0.653], P = 0.073, Fig 4), although this analysis was based on only two effect sizes.

**Fig 4:**
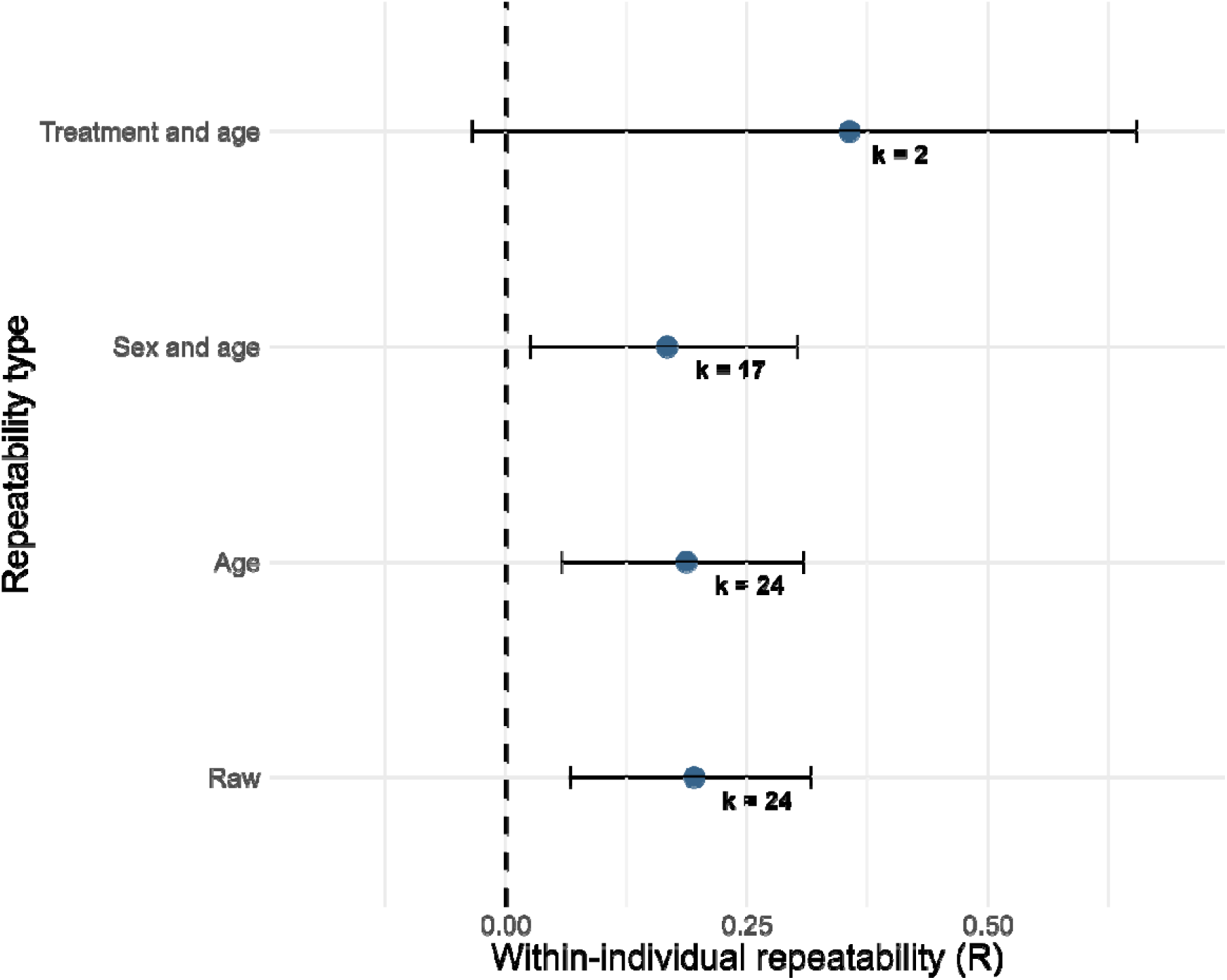
Within-individual repeatability of oxidative stress measures for the type of repeatability calculation. Repeatability type refers to what variables were controlled for in the model when estimating repeatability. Model estimates are shown along with the 95% confidence intervals. The dashed line indicates an R value of zero. The moderator is shown on the y axis, and R values (back transformed from Fishers Z values) are plotted on the x-axis.

Overall, these findings suggest that accounting for common covariates such as age and sex did not substantially alter repeatability estimates within the subset of studies analysed, although the limited number of available datasets restricts a broader inference regarding the drivers of intraindividual variation in oxidative stress.

### Are biomarkers of oxidative stress correlated?

Oxidative stress biomarkers are often assumed to reflect interrelated physiological processes, yet studies frequently report weak correlations among them (Cláudia Norte *et al*. 2008; Récapet *et al*. 2019). Our meta-analysis, showing generally low within-individual repeatability for oxidative stress markers further complicates this assumption. From a statistical perspective, low repeatability can substantially attenuate detectable correlations among biomarkers because high within-individual variation introduces additional unexplained variance into observed associations. Consequently, weak repeatability may contribute to generally weak correlations reported among oxidative stress markers.

Using individual-level raw data available from included studies, we assessed correlations between commonly measured oxidative stress biomarkers, including protein carbonyls – superoxide dismutase (carbonyl-SOD), malondialdehyde – antioxidant capacity (MDA-OXY), antioxidant capacity – superoxide dismutase (OXY-SOD), malondialdehyde – superoxide dismutase (MDA-SOD), and reactive oxygen metabolites – antioxidant capacity (dROMs-OXY) (Fig 5). Most pairwise correlations were weak and statistically non-significant (Table S7). However, we identified a modest positive correlation between dROMs and OXY (r = 0.250, 95% CI = [0.091, 0.396]; Table S7, Fig 4). This could suggest that both markers exhibit partially synchronous temporal variation within individuals despite relatively low repeatability estimates. Nevertheless, the overall pattern suggests that oxidative stress markers often exhibit weak or inconsistent correlations across studies and supports the use of longitudinal and multivariate approaches for investigating oxidative physiology in vivo.

**Fig 5:**
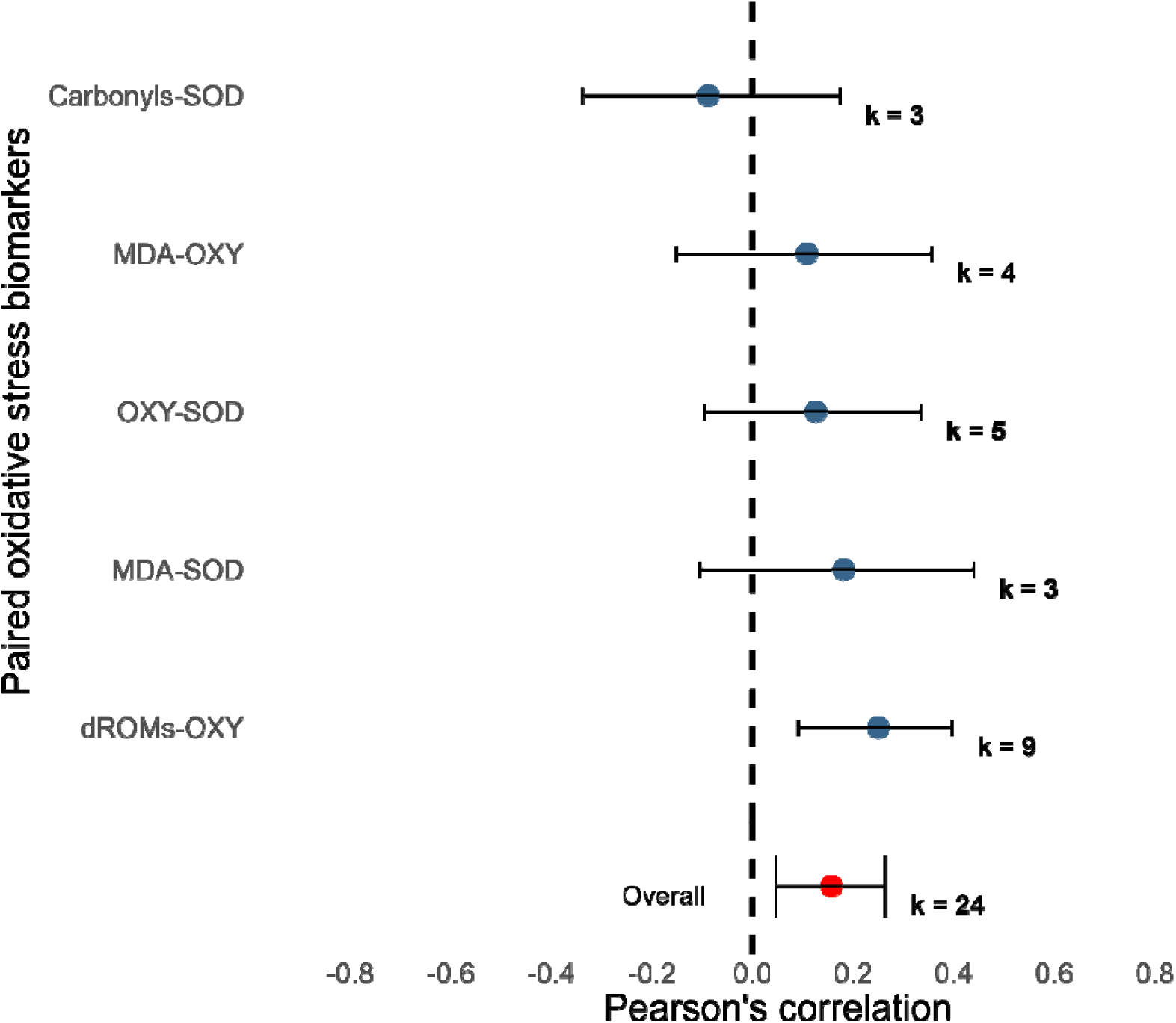
Pearson’s correlation for pairs of oxidative stress markers. The model estimates are shown along with 95% confidence intervals. The dashed line indicates where Pearson’s correlation is zero. The paired oxidative stress biomarkers are shown on the y axis and Pearson’s correlation (back transformed from Fishers Z) is shown on the x axis. The red point indicates the model estimate for the intercept-only model along with the 95% confidence intervals. “k” represents the number of effect sizes.

### Implications of low repeatability for statistical power and study design: A simulation-based case study on oxidative stress and telomere length

Our meta-analysis revealed that oxidative stress markers exhibit low within-individual repeatability, which reduces the ability to detect biologically meaningful associations between oxidative stress and other traits. To explore the potential implications of low repeatability for statistical inference, we conducted an individual-based simulation examining the relationship between oxidative stress and telomere length under three hypothetical biological scenarios: (1) oxidative stress fully contributing to telomere attrition, (2) moderate causality, and (3) weak causality. These simulations were designed as heuristic examples intended to illustrate how repeatability may influence statistical power under different assumptions, rather than to provide generalized recommendations for study design. All simulations were parameterized using biological values within the natural range of a hypothetical great tit population (see Supplementary Materials).

Our simulations suggested that under scenarios of low repeatability (ICC = 0.2), substantially larger sample sizes were often required to achieve sufficient statistical power compared to scenarios with moderate or high repeatability (Fig 6A). A previous meta-analysis examining oxidative stress and telomere length correlations found that only 16 studies included more than 100 individuals out of 37 studies (Wilbourn *et al*. 2018). Under scenarios assuming weaker relationships between oxidative stress and telomere attrition, statistical power declined further, particularly at smaller sample sizes (Fig 6A).

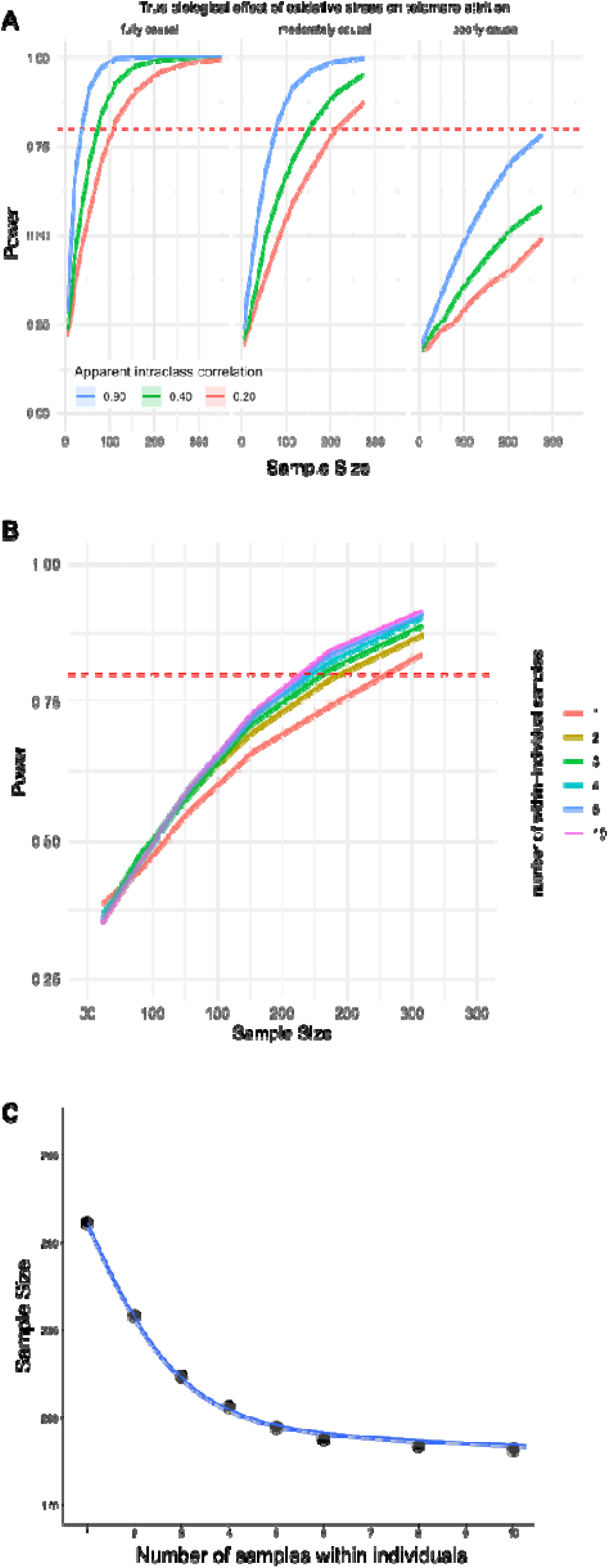

Depending on the study system, increasing the number of repeated measurements per individual may improve statistical power more efficiently than increasing total sample size alone. Our simulations indicated that increasing repeated oxidative stress measurements per individual reduces the total sample size required to achieve equivalent statistical power (Fig 6B). Within the moderate causality scenario, three to four repeated measurements per individual reduced required sample sizes by approximately 25-30%, beyond which additional sampling yielded diminishing returns. These simulations therefore illustrate how longitudinal sampling designs may partially offset the statistical consequences of low repeatability and may additionally create opportunity to investigate the sources of intraindividual variation in oxidative stress.

## Discussion

The utility of physiological biomarkers in ecology and evolutionary biology depends on their statistical accuracy in capturing biologically meaningful information, correlating individuals’ environmental contexts to fitness. Our meta-analysis demonstrates that oxidative stress markers exhibit low repeatability on average. Furthermore, the observed intraindividual variation appears largely independent of a range of predictor variables available that were to us for estimating conditional repeatability. The apparent high and unexplained intra-relative to inter-individual variation has important implications for the utility of oxidative stress measurements in ecological and physiological evolutionary studies because informative biomarkers that indicate variation in individual quality, environmental stress, life-history trade-offs, or fitness require at least moderate conditional repeatability. Importantly, our findings do not imply that oxidative stress markers lack biological relevance or cannot be useful indicators of physiological state in the case of disease or pathology. Rather, they suggest that low repeatability limits their statistical power to capture individual differences *in the healthy population*.

This distinction is particularly important as oxidative stress biomarkers are used differently across disciplines. In ecological and evolutionary studies, oxidative stress measurements are often interpreted in the context of variation in individual quality, physiological costs associated with reproduction, environmental stress, immune activation, survival, or ageing (A. Cohen *et al*. 2008; Alonso-Álvarez *et al*. 2010; Boonekamp *et al*. 2017; Herborn *et al*. 2016; Kosaruk *et al*. 2023; Pintus & Ros-Santaella 2021; Zhang *et al*. 2023). Such approaches implicitly assume that variation in oxidative stress captures among individual variation with sufficient accuracy. In contrast, biomedical and veterinary applications tend to interpret oxidative stress relative to physiological thresholds or disease states, where transient deviations from normal ranges may themselves be informative (Dhama *et al*. 2019; Giustarini *et al*. 2009). Reviews across biomedical and ecological disciplines have repeatedly highlighted that oxidative stress markers can behave dynamically depending on study system, organism, and environment (Beaulieu & Costantini 2014; Costantini 2019; Costantini *et al*. 2010). Our findings call for further research aimed at investigating the causes of intraindividual variation in the healthy population to increase the utility of oxidative stress measurements beyond their diagnostic use in the clinical context. The discussion below highlights several important details we found from our analyses that may offer a starting point to future research on the causes of intraindividual variation in oxidative stress.

Despite the low overall repeatability, several studies included in our meta-analysis reported relatively high repeatability of oxidative stress measurements (R > 0.6). What do these studies have in common that may explain their high repeatability? While it is not possible to make any definitive conclusions at this point, several tentative patterns emerged (Table S8). First, markers of lipid peroxidation, particularly MDA quantified using high-performance liquid chromatography (HPLC), frequently exhibited relatively high repeatability in both wild (Bodey *et al*. 2019; Christensen *et al*. 2015) and controlled settings (Orquera-Arguero *et al*. 2023), suggesting that some lipid peroxidation markers may retain stable among-individual differences across time. High and moderate repeatability was also observed for other biomarkers such as vitamin E and SOD under experimental conditions (Marasco *et al*. 2017; Velázquez-Cantón *et al*. 2018), as well as OXY, SOD, and peroxidation index in wild populations (Eikenaar *et al*. 2020; Récapet *et al*. 2019; Těšický *et al*. 2021).

High repeatability was also observed in some experimental or controlled environments, including supplementation studies and livestock systems, suggesting that reduced environmental heterogeneity or more stable physiological conditions may increase temporal consistency. Conversely, in wild populations, higher repeatability tended to occur under prolonged or relatively predictable physiological challenges such as migration, lactation, or overwinter survival (Bodey *et al*. 2019; Christensen *et al*. 2016; Eikenaar *et al*. 2020; Orquera-Arguero *et al*. 2023). Together, these observations suggest that repeatable oxidative phenotypes may emerge under sustained extreme metabolic or ecological conditions. However, these patterns were not universally consistent across studies or biomarkers, and even markers exhibiting relatively high repeatability in some contexts often showed weak repeatability elsewhere. Consequently, despite some markers performing better than others, we failed to identify broadly generalisable contexts in which oxidative stress markers consistently exhibited repeatability levels that would support strong ecological inference about fitness or long-term physiological state. It is important to recognise that studies included in our meta-analysis did not select biomarkers at random. Researchers generally chose assays based on the biology of their study system, the physiological process of interest, and current understanding of oxidative physiology. Consequently, if repeatability remains low across a diverse literature, it suggests that low repeatability may not simply reflect inappropriate biomarker choice. Instead, it may reflect a more fundamental feature of oxidative physiology itself, namely that oxidative stress is highly dynamic and context dependent, even when measured using markers that are considered relevant to the study system.

These findings further emphasis that oxidative stress markers should not be treated as a homogeneous biomarker class. Oxidative stress encompasses a wide range of biologically distinct molecules, including reactive oxygen species, enzymatic antioxidants, non-enzymatic antioxidants, lipid peroxidation products, protein oxidation products, and DNA damage markers. Some of the products we can measure are derived from intra-cellular processes, others are extracellular. This repertoire of molecules and enzymes differ substantially in biochemical function, half-life, tissue specificity, temporal stability, and environmental responsiveness (Costantini 2019; Costantini *et al*. 2010; Pamplona & Costantini 2011). Previous work has similarly shown that oxidative stress markers often exhibit weak or inconsistent covariation within individuals (A. Cohen *et al*. 2008). Grouping these biomarkers together by supposed function or class therefore represents a simplification that may obscure important nuances in the complex physiology of different aspects of oxidative stress.

Nevertheless, further subdivision of biomarkers within our meta-analysis would have resulted in over-parametrisation and insufficient sample sizes for many categories. Our results therefore highlight the need for future studies to adopt a multi-biomarker approach to gain more insight into the different aspects of oxidative stress physiology.

A key challenge when interpreting low repeatability is distinguishing between biological variability and technical measurement error. High variation in oxidative stress markers may arise not only from biological plasticity, but also from assay noise, storage effects, batch effects, calibration instability, or other sources of methodological variation (Aravapally *et al*. 2025; Miranda *et al*. 2026). Although we were unable to explicitly partition technical and biological variance components within the present meta-analysis, several studies included within the meta-analysis reported high technical repeatability or low inter-assay coefficients of variance (CV) despite low biological repeatability estimates (Table S1). Across studies reporting assay precision, technical repeatability was generally high, in the region of 69-98%, whereas inter- and intra-assay CVs were commonly below 10%. These observations suggest that assay imprecision due to technical flaws alone does not suffice to explain the general low repeatability estimates observed across studies. This points to substantial within-individual biological variation and environmentally driven variability as important contributors to oxidative stress dynamics.

The temporal scale of oxidative stress variation is also likely to be important. Even when oxidative stress genuinely influences fitness or ageing, rapid fluctuations in oxidative physiology may obscure detectable associations if measurements are obtained at biologically short timescales. At present, we are not aware of dedicated studies examining fine scaled temporal patterns of within-individual variability of oxidative stress through the life course in free living organisms. However, experimental studies in cell cultures have demonstrated highly dynamic oxidative fluctuations even under controlled conditions (Ayuso *et al*. 2020; Halliwell 2003; Musakhanian *et al*. 2022; Shiva Shankar Reddy *et al*. 2007). This does not mean per se that even greater variability is expected in vivo as a complex system may offer greater capacity to buffer against extrinsic and intrinsic factors, offering greater stability. Among the studies in our meta-analysis that collected repeated measurements over relatively short timescales (e.g. one month or less), (Cram *et al*. 2015; Eikenaar *et al*. 2020; Gormally *et al*. 2019; Kolling *et al*. 2022; Noguera *et al*. 2017; Orquera-Arguero *et al*. 2023; Sauerwein *et al*. 2020; Schull *et al*. 2016; Velázquez-Cantón *et al*. 2018; Viblanc *et al*. 2018) most reported low repeatability, although a minority of studies under controlled conditions or with multiple repeated measurements detected moderate repeatability (Eikenaar *et al*. 2020; Orquera-Arguero *et al*. 2023; Velázquez-Cantón *et al*. 2018). These findings suggest that oxidative stress may often behave as a highly plastic physiological system responding to environmental and physiological conditions. To better characterise and understand the causes of this variation, we need longitudinal studies that collect repeated measurements at a granular timescale and preferably throughout the life course. This would enable identifying the drivers of intraindividual variation in oxidative stress potentially improving the utility of oxidative stress measurements as a biomarker of body condition and fitness.

Our analysis additionally revealed generally weak correlations among oxidative stress markers, reinforcing previous findings that oxidative physiology is multivariate (Christensen *et al*. 2015; Mallard *et al*. 2020). These findings presents a conceptual challenge because oxidative stress markers are often interpreted as reflecting interconnecting physiological pathways (Costantini *et al*. 2010; Skrip & McWilliams 2016; Smith *et al*. 2016). However, weak correlations may emerge naturally when biomarkers exhibit low repeatability and fluctuate independently throughout time. Under these conditions, high within-individual variance can substantially attenuate observed correlations among traits. Importantly, low repeatability does not preclude strong correlations if multiple traits fluctuate synchronously in response to shared physiological processes or environmental drivers. We found some evidence for this possibility in the positive correlation observed between ROMs and OXY. This suggests that some oxidative stress markers may exhibit coordinated temporal dynamics despite relatively low individual repeatability.

In conclusion, our study highlights the importance of understanding the ecological, physiological, and methodological drivers of intraindividual variation. Future work should prioritise longitudinal study designs and carefully select biomarkers depending on the question and study system. We also advocate more crosstalk among the fields of ecology, veterinary medicine, and biomedicine to better understand under what conditions and criteria oxidative stress biomarkers can provide the most insight into physiological state, ageing, and fitness.

## Availability of data and materials

The code and data used in this study will be made available online following peer review at the time when the manuscript is accepted for publication.

## Acknowledgments

Thanks to all of the authors who responded to our data enquiries including Quentin Schull, Sarah-Guindre-Parker, Helga Sauerwein, Melissah Rowe, Brenna Gormally, Aurora Ramierez-Perez, Mireia Blanco, Shelia Stivanin, Valeria Marasco, and Martin Tesicky.

## Author contributions

RR, DMD, JB conceived and designed the study. RR and JB developed the simulation. RR ran the meta-analysis and drafted the manuscript with help from DMD and JB. All authors read and approved the final version.

