## Supplementary materials for "Oxidative stress markers have low repeatability: A meta-analysis and simulation study with implications for measuring physiological condition and fitness"

**S1 text - Individual-based simulation methods.**

We developed an individual-based simulation in R to illustrate the potential statistical consequences of low repeatability for detecting associations between oxidative stress and telomere length. The simulation was intended as a heuristic and illustrative framework rather than a predictive model for study design or a biologically realistic representation of telomere dynamics. We chose telomere length because oxidative stress is frequently hypothesised to contribute to telomere attrition, yet empirical correlations between oxidative stress and telomere length are often weak and inconsistent across studies.

The simulations explored how statistical power changes under different scenarios of oxidative stress repeatability and varying assumptions regarding the extent to which oxidative stress contributes to telomere attrition. Importantly, the simulation does not assume that oxidative stress is the sole or primary determinant of telomere loss. Instead, the different causality scenarios were included to illustrate how the detectability of relationships changes under varying assumptions regarding biological effect size.

To provide biologically plausible parameter ranges, we used starting values for telomere length, oxidative stress, and mortality that roughly reflected a hypothetical population of great tit (*Parus major*) fledglings. Each simulation generated a population with a normal distribution of telomere length (mean = 30 kb, SD = 8 kb) and oxidative stress (mean = 1500, SD = 450). We then simulated annual survival, oxidative stress dynamics, and telomere loss throughout life. Annual mortality was simulated using a Gompertz survival function (Baseline mortality rate = 0.02, age-dependent increase = 0.15) to emulate realistic age-dependent mortality from non-specific causes. Individuals additionally died if telomere length fell below a critical threshold (<1000 bp), although most mortality arose from the Gompertz survival function rather than telomere attrition itself.

Each year an individual survived, oxidative damage was simulated using the following equation:

Damage = OS_birth_ + ƴ * (1-r)

Where r represents repeatability and ƴ is a random scaling value ranging between -1500 and 4500. This parameter range was selected to generate substantial within-individual variation and was not intended to represent empirically derived biological limits. When repeatability approached 1, oxidative damage was largely determined by oxidative stress at fledging, whereas lower repeatability values generated increasingly stochastic within-individual variation over time.

Importantly, the repeatability parameter in the simulation does not directly correspond to the resulting intraclass correlation coefficient (ICC). Consequently, we calculated the apparent ICC values emerging from the simulated datasets and used these realized ICC estimates for subsequent power analysis. Negative damage values were permitted in a small proportion of cases to reflect occasional telomere elongation observed empirically in some populations. Including or excluding negative damage values had negligible effects on the resulting power estimates.

We simulated repeatability scenarios corresponding approximately to apparent ICC values of 0.2, 0.4, and 0.9. The ICC = 0.2 scenario reflected the approximate mean repeatability observed in our empirical meta-analysis, whereas higher ICC scenarios represented progressively more stable oxidative stress phenotypes.

Telomere loss was generated annually as a function of oxidative stress and cell division using the following equation:

TL_t+1_ = TL_t_ – damage - cell division

Where TL represents telomere length at age t, damage is as defined above, and cell division is a random variable drawn from a normal distribution (mean = 100, SD = 20). The contribution of cell division was comparatively small because cell turnover is highest during early development and decreases substantially after fledging. We simulated three causality scenarios representing weak (30%), intermediate (60%), and strong (100%) contributions of oxidative stress to telomere attrition. Even in the fully causal scenario, some telomere loss remained attributable to cell division. These scenarios should not be interpreted as empirically validated biological effect sizes but rather as illustrative examples intended to explore how repeatability influences statistical detectability under varying assumptions.

Each simulated population generated longitudinal trajectories of oxidative stress and telomere length. We then calculated the correlation between oxidative stress and telomere length at median survival age of each simulated population. Simulations were repeated across combinations of sample size (n = 10-350), repeatability scenario (ICC = 0.2, 0.4, and 0.9), and causalty scenario (30%, 60%, and 100%). For each parameter combination, simulations were repeated 1000 times to estimate statistical power and generate power-sample size curves with 95% confidence intervals.

To illustrate how repeatability and causality influence the apparent relationship between oxidative stress and telomere length, we additionally plotted individual-level trajectories from a subset of simulations between ages 5.5 and 6.5 years (Fig S1).

Finally, we investigated the impact of collecting repeated oxidative stress measurements within individuals on statistical power. Additional simulation scenarios varied the number of oxidative stress measurements collected per individual sampled from 1-10 repeated samples. These analyses were intended to illustrate how repeated sampling may improve statistical power under low repeatability conditions rather than provide universal recommendations for study design.

Each population simulated, generated longitudinal data on within-individual oxidative stress and telomere trajectories. We then determined the correlation between oxidative stress and telomere length at the median survival time of each population. This process was repeated for different combinations of population sample sizes (ranging from n=10 to n=350), repeatability scenarios (ICC= 20%, 40%, and 90%), and causality scenarios (30%, 60%, and 100%). For each scenario, we computed the statistical power, and scenarios were repeated 1000 times to obtain precise power-sample size curves and 95% confidence intervals. To illustrate how repeatability and causality influence the apparent relationship between oxidative stress and telomere length, we plotted individual-level data from a subset of simulations at ages 5.5-6.5 years (Figure S1).

Finally, we investigated the impact of collecting repeated oxidative stress measurements within individuals on the statistical power for detecting a relationship between oxidative stress and telomere length. To this end we added a few additional scenarios regarding the number of oxidative stress measurements taken from each individual (ranging from 1-10).

The simulation code used in this study is available on Github (<https://github.com/RachelRReid/Oxidative-stress-simulation>) and archived on Zenodo (<https://doi.org/10.5281/zenodo.16760050>).

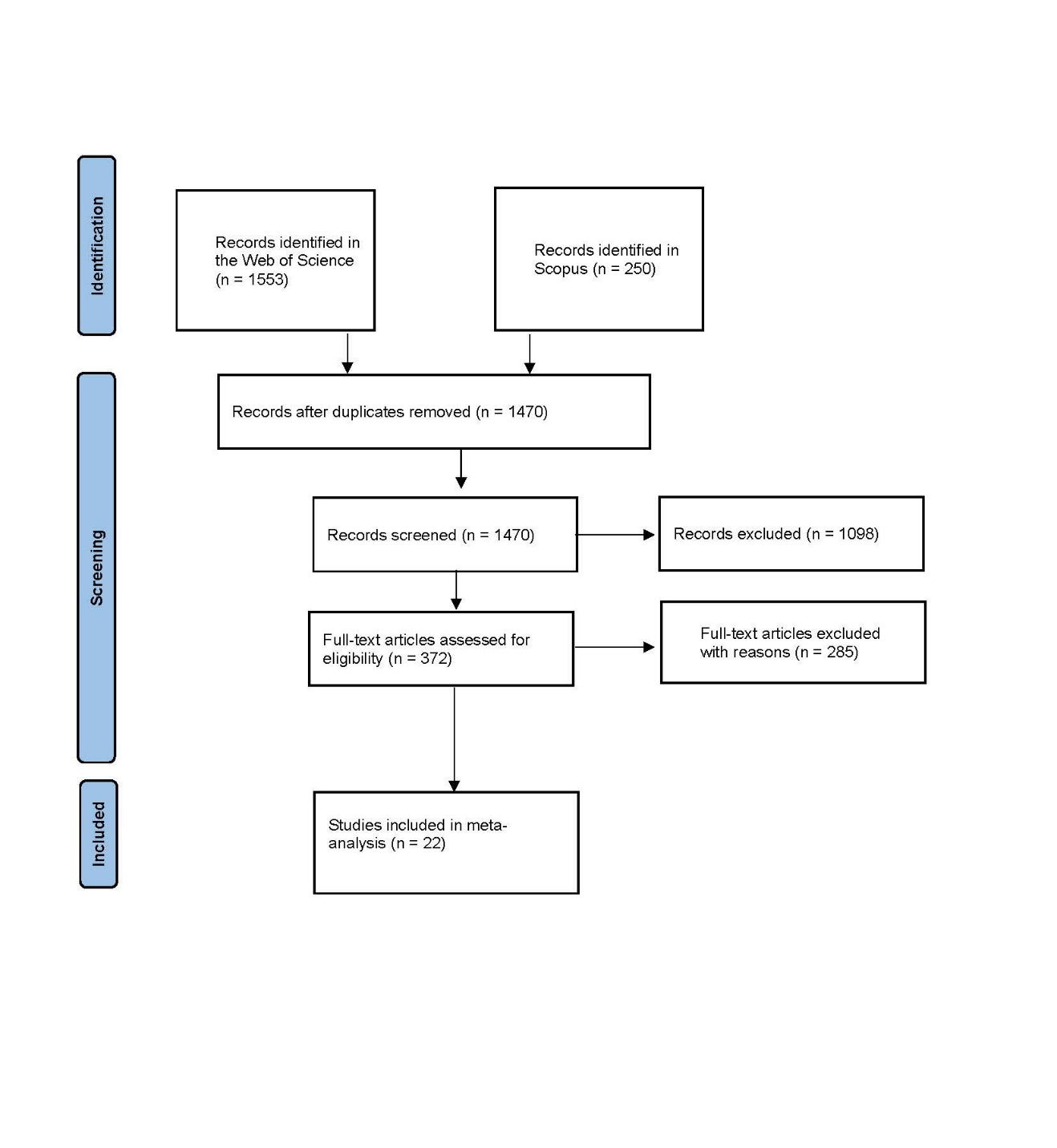

Figure S1. PRISMA diagram showing each stage of the literature screening process.

**Table S1.** Information about technical repeatability of the assays in each study if reported alongside the ICC calculated.

| Study ID | Marker | Technical repeatability/CV | Biological ICC |
| --- | --- | --- | --- |
| Arguero2023 | MDA | Inter assay CV 7.3%  Intra assay CV 4.6% | 0.93 |
| Christenson2015 | Protein carbonyls | Inter assay CV 8.5% | <0 |
| Christenson2015 | MDA | Inter assay repeatability 0.91 | 0.08-0.7 |
| Christenson2015 | OXY | Inter assay CV 3% | 0.04-0.06 |
| Christenson2015 | SOD | Inter assay CV 3.7% | <0 |
| Cram2014 | OXY | Intra assay repeatability R = 0.88 | 0.25-0.39 |
| Cram2014 | SOD | Intra assay repeatability R = 0.72 | 0.14-0.35 |
| Cram2014 | MDA | Intra assay repeatability R = 0.78 | 0-0.4 |
| Cram2014 | Uric acid | Intra assay repeatability R = 0.79 | 0.06-0.48 |
| Gormally2019 | Uric acid | Inter assay CV 2.3%  Intra assay CV 2.5% | 0-0.14 |
| Hau2015 | ROMs | Inter assay CV 7.4%  Intra assay CV 5.8% | 0.17-0.46 |
| Hau2015 | OXY | Inter assay CV 6.7%  Intra assay CV 6% | 0-0.22 |
| Hau2015 | GPX | Inter assay CV 8%  Intra assay CV 5.6% | 0-0.23 |
| Herborn2015 | ROMs | Inter assay CV 4.7%  Intra assay CV 10.2% | 0.03-0.06 |
| Herborn2015 | OXY | Inter assay CV 8.3%  Intra assay CV 9.9% | <0 |
| Marasco2017 | 8OHdG | Inter assay CV 4.8%  Intra assay CV 7.9% | 0-0.12 |
| Marasco2017 | OXY | Inter assay CV 3.6%  Intra assay CV 4.2% | 0-0.18 |
| Marasco2017 | SOD | Inter assay CV 3.6%  Intra assay CV 8.3% | 0.33-0.59 |
| Marasco2017 | Protein carbonyls | Inter assay CV 8.9%  Intra assay CV 6% | <0 |
| Nogura2017 | GPX | Intra assay repeatability R = 0.97 | 0-0.03 |
| Nogura2017 | MDA | Intra assay repeatability R = 0.86 | 0.001-0.09 |
| Nogura2017 | Uric acid | Intra assay repeatability R = 0.98 | 0.03-0.09 |
| Parker2013 | ROMs | Intra assay CV 8.5% | 0.27 |
| Parker2013 | OXY | Intra assay CV 3.4% | 0 |
| Parker2018 | ROMs | Inter assay CV 10.4%  Intra assay CV 5.9% | 0.11-0.44 |
| Parker2018 | OXY | Inter assay CV 11.9%  Intra assay CV 5.4% | 0-0.27 |
| Recapet2019 | OXY | Inter assay repeatability R = 0.69  Intra assay repeatability R = 0.8 | 0-0.46 |
| Recapet2019 | ROMs | Inter assay repeatability R = 0.86  Intra assay repeatability R = 0.91 | <0 |
| Sauerwein2020 | ROMs | Inter assay CV 9.6%  Intra assay CV 1.7% | 0.036-0.35 |
| Sauerwein2020 | OXY | Inter assay CV 0.5%  Intra assay CV 1.1% | 0.12-0.29 |
| Sauerwein2020 | TBARs | Inter assay CV 8.3%  Intra assay CV 5.4% | 0.34-0.55 |
| Sauerwein2020 | AOPP | Inter assay CV 2.9%  Intra assay CV 1.9% | 0.22-0.37 |
| Schull2016 | ROMs | Inter assay CV 5.4%  Intra assay CV 5.3% | 0.069 |
| Schull2016 | OXY | Intra assay CV 4.2%  Inter assay CV 3.3% | 0.19 |
| Schull2016 | SOD | Inter assay CV 7%  Intra assay CV 3.1% | 0.09 |
| Tesicky2021 | TOBR | Intra assay CV 8.9% | <0 |
| Tesicky2021 | Thiols | Inter assay CV 15%  Intra assay CV 7.7% | 0-0.27 |
| Tesicky2021 | SOD | Inter assay CV 7.5%  Intra assay CV 29.2% | 0.47-0.67 |
| Tesicky2021 | GPX | Inter assay CV 21%  Intra assay CV 7.6% | 0.38-0.49 |
| Viblanc2017 | ROMs | Inter assay CV 13.9%  Intra assay CV 3.7% | 0.1 |
| Viblanc2017 | OXY | Inter assay CV 7.1%  Intra assay CV 4.7% | 0.18 |

**Table S2.** List of oxidative stress biomarkers used in the meta-analysis alongside the type of damage or defence, the corresponding sample sizes are also listed.

| Measure type – full name | Acronym | Overall Biomarker | Type of damage or defence | Sample size |
| --- | --- | --- | --- | --- |
| 8-hydroxyl-2’-deoxyguanosine | 8OHdG | Oxidative damage | DNA damage | 4 |
| Advanced Oxidation Protein Products | AOPP | Oxidative damage | Protein damage | 2 |
| Malondialdehyde | MDA | Oxidative damage | Lipid peroxidation | 12 |
| Peroxidation index | Peroxidation index | Oxidative damage | Lipid peroxidation | 2 |
| Protein carbonyls | Protein carbonyls | Oxidative damage | Protein damage | 6 |
| Reactive Oxygen Metabolites | ROMs | Oxidative damage | Widespread damage | 19 |
| Thiobarbituric aid reactive substances | TBARs | Oxidative damage | Lipid peroxidation | 5 |
| Total oxidative burst responsiveness | TOBR | Oxidative damage | Widespread damage | 2 |
| Catalase | Catalase | Antioxidant defence | Enzymatic defence | 2 |
| Glutathione peroxidase | GPX | Antioxidant defence | Enzymatic defence | 8 |
| Reduced glutathione | GSH | Antioxidant defence | Non-enzymatic defence | 2 |
| Total antioxidant capacity of plasma | OXY | Antioxidant defence | Non-enzymatic defence | 30 |
| Plasma carotenoids | Plasma carotenoids | Antioxidant defence | Non-enzymatic defence | 4 |
| Superoxide dimutase | SOD | Antioxidant defence | Enzymatic defence | 13 |
| Thiols | Thiols | Antioxidant defence | Non-enzymatic defence | 2 |
| Uric acid | UA | Antioxidant defence | Non-enzymatic defence | 8 |
| Vitamin E | Vitamin E | Antioxidant defence | Non-enzymatic defence | 2 |

**Table S3.** List of authors that sent their raw data to be used in the meta-analysis when requested. The table also lists the title of the study that corresponded to the data as well as the journal it was published in and the year of publication.

| Corresponding author | Title of paper | Journal | Publication year |
| --- | --- | --- | --- |
| Quentin Schull | The oxidative debt of fasting: evidence for short- to medium-term costs of advanced fasting in adult king penguins | Journal of Experimental Biology | 2016 |
| Sarah Guindre-Parker | The oxidative costs of territory quality and offspring provisioning | Journal of Evolutionary Biology | 2013 |
| Sarah Guindre-Parker | No short-term physiological costs of offspring care in a cooperatively breeding bird | Journal of Experimental Biology | 2018 |
| Helga Sauerwein | Acute phase proteins and markers of oxidative status in water buffalos during the transition from late pregnancy to early lactation | Veterinary Immunology and Immunopathology | 2020 |
| Melissah Rowe | Exploratory behavior is associated with plasma carotenoid accumulation in two congeneric species of waterfowl | Behavioural Processes | 2015 |
| Brenna Gormally | Recovery from repeated stressors: Physiology and behavior are affected on different timescales in house sparrows | General and Comparative Endocrinology | 2019 |
| Aurora Ramierez-Perez | Effect of Selenium and Vitamin E Supplementation on Lactate, Cortisol, and Malondialdehyde in Horses Undergoing Moderate Exercise in a Polluted Environment | Journal of Equine Veterinary Science | 2018 |
| Mireia Blanco | Beef cows' performance and metabolic response to short nutritional challenges in different months of lactation | Research in Veterinary Science | 2023 |
| Shelia Stivanin | Milk production and haematological and antioxidant profiles of dairy cows supplemented with oregano and green tea extracts as feed additives | Brazilion Journal of Animal Science | 2022 |
| Valeria Marasco | Environmental conditions can modulate the links among oxidative stress, age, and longevity | Mechanisms of Ageing and Development | 2017 |
| Martin Tesicky | Longitudinal evidence for immunosenescence and inflammaging in free-living great tits | Experimental Gerontology | 2017 |

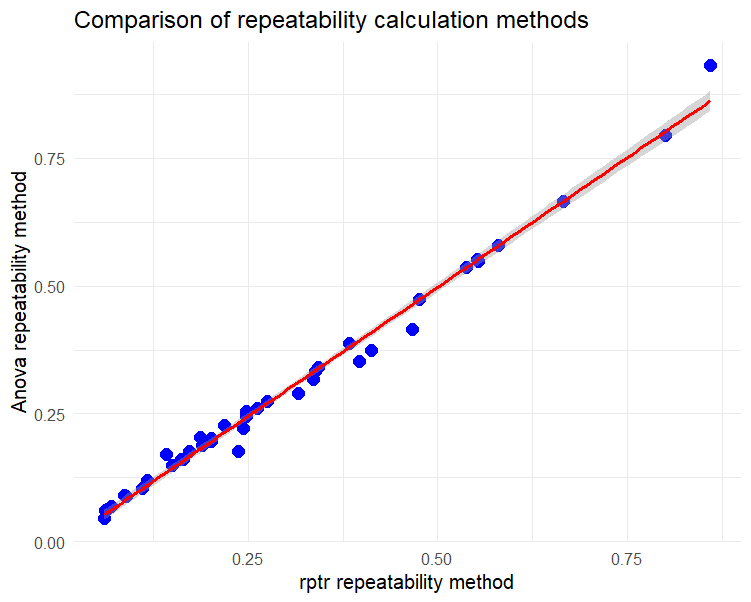

Figure S2. Correlation between the LMM and ANOVA methods of calculating repeatability. The blue points show the repeatability values and the red regression lines represent the correlation between the two methods, the shaded area around the line represents the 95% confidence intervals.

**Table S4.** List of the study species used in the meta-analysis alongside the type of environment the study took place in alongside the corresponding sample size.

| Study species Latin name | Common name | Taxa | Environment | Sample size |
| --- | --- | --- | --- | --- |
| Anas acuta | Northern Pintail | Aves | Captive | 2 |
| Anas platyrhynchos | Mallard Duck | Aves | Captive | 2 |
| Aptenodytes patagonicus | King Penguin | Aves | Wild Captive | 4 |
| Bos Taurus | Cattle | Mammal | Livestock | 11 |
| Branta bernicla | Light bellied Brent Geese | Aves | Wild | 8 |
| Bubalus bubalus | Buffalo | Mammal | Livestock | 8 |
| Equus caballus | Horse | Mammal | Livestock | 4 |
| Ficedula albicollis | Collared Flycatcher | Aves | Wild | 8 |
| Lamprotornis nitens | Glossy Starling | Aves | Wild | 2 |
| Lamprotornis superbus | Superb Starling | Aves | Wild | 4 |
| Larus michahellis | Yellow-legged Gull | Aves | Wild | 6 |
| Mungos mungo | Banded Mongoose | Mammal | Wild | 1 |
| Oenanthe oenanthe | Wheatear | Aves | Wild Captive | 6 |
| Ovis aries | Sheep | Mammal | Livestock | 8 |
| Parus major | Great tit | Aves | Wild | 10 |
| Passer domesticus | House Sparrow | Aves | Wild Captive | 2 |
| Phalacrocorax aristotelis | Shag | Aves | Wild | 4 |
| Plectrophenax nivalis | Snow Bunting | Aves | Wild | 2 |
| Plocepasser mahali | White-browed Sparrow Weaver | Aves | Wild | 8 |
| Taeniopygia castanotis | Zebra Finches | Aves | Captive | 15 |
| Turdus merula | Blackbird | Aves | Wild Captive | 6 |
| Urocitellus columbianus | Columbian Ground Squirrel | Mammal | Wild | 2 |

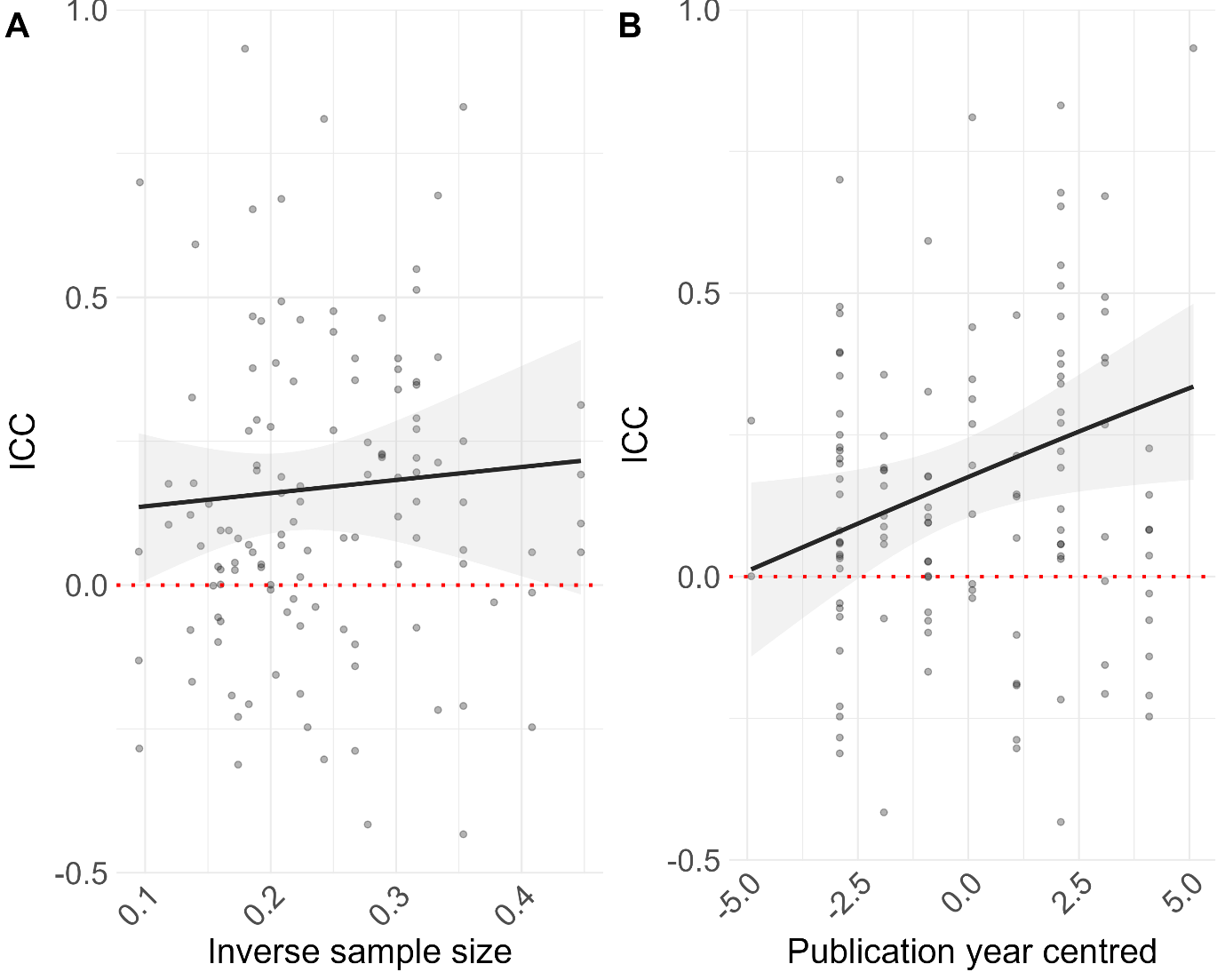

Figure S3. Scatter plot showing evidence of publication bias in the main dataset: (A) shows the relationship between the study sample size (this has been transformed using the square root of the inverse of the sample size) and ICC. (B) shows the relationship between publication year (this has been mean centered) and ICC. The solid line represents the model estimate and the shaded area around the line represents the 95% confidence intervals, the individual effect sizes are also shown.

**Table S5.** This table shows the model outputs for all moderators used on the full dataset. Bold estimates indicate confidence intervals (CI) that do not overlap zero. k is the number of effect sizes per level, and n is the number of studies. The values in the table have been back transformed from Fishers Z to ICC.

|  | Estimate [95%CI] | k | n |
| --- | --- | --- | --- |
| *Intercept only model*  *I2 total= 57.399*  *I2 Obs ID= 51.295*  *I2 Study= 6.104* | **0.164 [0.093, 0.233]** | 123 | 22 |
| Log average time between measures (days) | 0.022 [0.006, 0.038] | 65 | 21 |
| Log average number of samples per individual | **0.097 [0.064, 0.130]** | 68 | 22 |
| Taxa  *Aves*  *Mammal* | **0.155 [0.062, 0.245]**  **0.212 [0.058, 0.356]** | 89  34 | 15  7 |
| Environment  *Livestock*  *Wild*  *Captive*  *Wild captive* | **0.391 [0.178, 0.569]**  0.119 [-0.003, 0.239]  0.158 [-0.07, 0.371]  0.165 [-0.052, 0.367] | 23  63  19  18 | 4  11  3  4 |
| Overall Biomarker  *Antioxidant defence*  *Oxidative damage* | **0.171 [0.082, 0.258]**  **0.156 [0.056, 0.253]** | 71  52 | 20  20 |
| Study Type  *Manipulation*  *Non-manipulation* | **0.208 [0.091, 0.319]**  **0.146 [0.053, 0.230]** | 40  83 | 12  18 |
| Sex  *Female*  *Male* | 0.167 [0.065, 0.266]  0.154 [0.03, 0.273] | 62  37 | 16  12 |

**Table S6.** This table shows the model output for the subgroup analysis on both “Oxidative damage” and “Antioxidant defence” subgroups. Bold estimates indicate confidence intervals (CI) that do not overlap zero. k is the number of effect sizes per level, and n is the number of studies. The values in the table have been back transformed from Fishers Z to ICC.

|  | Estimate [95%CI] | k | n |
| --- | --- | --- | --- |
| Type of oxidative damage  *DNA damage*  *Protein damage*  *Lipid peroxidation*  *Widespread* | 0.135 [-0.224, 0.461]  -0.068 [-0.346, 0.220]  **0.368 [0.182, 0.529]**  0.059 [-0.137, 0.252] | 4  8  19  21 | 2  4  10  10 |
| Type of antioxidant defence  *Enzymatic*  *Non-enzymatic* | **0.204 [0.081, 0.320]**  **0.136 [0.042, 0.228]** | 23  48 | 9  20 |

**Table S7.** This table shows the model output for Pearson’s correlation between paired oxidative stress biomarkers. Bold estimates indicate confidence intervals (CI) that do not overlap zero. k is the number of effect sizes per level, and n is the number of studies. The values in the table have been back transformed from Fishers Z to Pearsons correlation.

|  | Estimate [95%CI] | k | n |
| --- | --- | --- | --- |
| *Intercept only model*  *I2 total= 87.298*  *I2 Obs ID= 73.026*  *I2 Study= 14.272* | **0.157 [0.046, 0.264]** | 24 | 14 |
| Oxidative stress pair  *Protein Carbonyls – SOD*  *dROMs-OXY*  *MDA-OXY*  *MDA-SOD*  *OXY-SOD* | -0.088 [-0.338, 0.173]  **0.250 [0.091, 0.396]**  0.108 [-0.153, 0.355]  0.181 [-0.105, 0.439]  0.126 [-0.095, 0.335] | 3  9  4  3  5 | 3  8  4  3  5 |

**Table S8:** Study level information corresponding to any moderate or high (>0.5) ICC estimates calculated.

| Study ID | Species | Biomarker | ICC | Assay type | Environment | Sampling interval | No repeats | Sex | Lifestage | Sample size | Manipulation |
| --- | --- | --- | --- | --- | --- | --- | --- | --- | --- | --- | --- |
| Arguero2023 | Cattle | MDA | 0.93 | HPLC | Livestock | 3 days | 3 | F | Adult | 31 | NA |
| Bodey2020 | Light Bellied Brent Geese | MDA | 0.68 | HPLC | Wild | Unknown | 2 | M | Adult | 9 | NA |
| Bodey2020 | Light Bellied Brent Geese | MDA | 0.51 | HPLC | Wild | Unknown | 2 | F | Adult | 10 | NA |
| Bodey2020 | Light Bellied Brent Geese | OXY | 0.83 | Cayman Colorimetric test | Wild | Unknown | 2 | M | Adult | 8 | NA |
| Canton2018 | Horses | Vitamin E | 0.81 | HPLC | Livestock | 7 days | 12 | Both | Adult | 17 | Supplemented feed |
| Christenson2015 | Sheep | MDA | 0.7 | HPLC | Wild | 1 year | 2 | F | Adult | 109 | NA |
| Cram2014 | White Browed Weaver | Uric acid | 0.48 | Cayman assay | Wild (breeding) | 30 days | 2 | Both | Adult | 16 | NA |
| Hau2015 | Blackbird | ROMs | 0.46 | dROMs assay | Wild captive | 1 year | 2 | Both | Adult | 12 | Regime of stressor exposure |
| Recapet2019 | Collared Flycatcher | OXY | 0.46 | Oxy adsorbent assay | Wild (breeding) | 1 year | 2 | M | Adult | 20 | NA |
| Sauerwein2020 | Buffalo | TBARs | 0.55 | TBARs test | Livestock (breeding) | 15 days | 4 | F | Adult | 10 | NA |
| Tesicky2021 | Great tit | SOD | 0.47 | Kinetic assay | Wild (breeding) | 365 days | 2 | M | Adult | 29 | NA |
| Tesicky2021 | Great tit | GPX | 0.49 | GPX cellular activity assay kit | Wild (breeding) | 365 days | 2 | F | Adult | 23 | NA |
